# Employing Microsoft Sway as a Repository for Data Generated in Pharmaceutics Experimental Courses: A Case study on Topical Formulation Characterization

**DOI:** 10.1101/2024.11.25.620827

**Authors:** Qingxin Chen, Rui Luo, Xin Pan, Chuanbin Wu, Zhengwei Huang

**Affiliations:** College of Pharmacy, Jinan University, Guangzhou 511443, Guangdong, PR China; State Key Laboratory of Bioactive Molecules and Druggability Assessment, Guangdong Basic Research Center of Excellence for Natural Bioactive Molecules and Discovery of Innovative Drugs, College of Pharmacy, Jinan University, Guangzhou 511443, Guangdong, PR China; School of Pharmaceutical Sciences, Sun Yat-Sen University, Guangzhou 510006, China

**Keywords:** Microsoft Sway, Data Repository, Undergraduate, Pharmaceutics Experimental Course

## Abstract

Microsoft Sway is an easy-to-use online creative software developed by Microsoft. It has many advantages as follows: easy to operate and does not require long-term practice and learning; network links can be created and accessed by devices such as Personal Computer (PC) and mobile terminals; Educators and Learners are allowed to browse this repository at anytime to facilitate the application prospects of mining relevant data. Therefore, the author proposes that Microsoft Sway may be a very promising data repository for pharmaceutical experimental teaching courses, and is expected to be extended to undergraduate experimental teaching in other natural science fields.

## Introduction

In undergraduate pharmacy education of China and many other countries, there are four main courses, namely medicinal chemistry, pharmacology, pharmaceutical analysis and pharmaceutics. The teaching of these four courses is typically delivered with theoretical and experimental curricula. Taking the example of pharmaceutics, theoretical and experimental courses are set. The main teaching content of pharmaceutics experimental courses usually includes but is not limited to the drawing of ternary phase diagrams, the determination of critical micelle concentration (CMC) of surfactants, the exploration of solubilization of solubilizers, the fluidity test of powders, and the preparation and evaluation of suspensions, emulsions, injections, tablets and ointments, *etc*.

During the training process of pharmaceutics experimental courses, a large amount of experimental data is often generated, such as the CMC of sodium lauryl sulphate, the dissolution rate of aspirin tablets, as well as the release rate of salicylic acid topical ointments with different matrices. In the big-data era, “there is no useless data” has been increasingly accepted by various disciplines(1). Thus, we are reminded that the experimental data generated in the pharmaceutics experimental course may also have potential application value. For example, it may provide data support for the development of related scientific research topics and teaching reforms in the future. In such a context, we should consider the systematic collection, collation and analysis of data generated in pharmaceutics experimental courses.

Drawing on the author’s experience of teaching and researching, we propose that Microsoft Sway can be a very promising repository for data generated in pharmaceutics experimental courses. Microsoft Sway is a Microsoft Office application that can be used to easily create and share interactive reports, narratives and presentations. It is free for anyone with a Microsoft account. It is worth mentioning that, being owned by Microsoft, this software can generate works based on the commonly used Microsoft PowerPoint documents (.ppt or .pptx), thus very convenient. There have been a series of studies that have applied it to teaching research and reform. In previously published studies, educators primarily use Microsoft Sway to optimize self-directed learning modes, to propose pedagogies under COVID-19 pandemic circumstances, and other aspects. Relevant information from classic studies is summarized in **Table 1**.

**Table 1.**
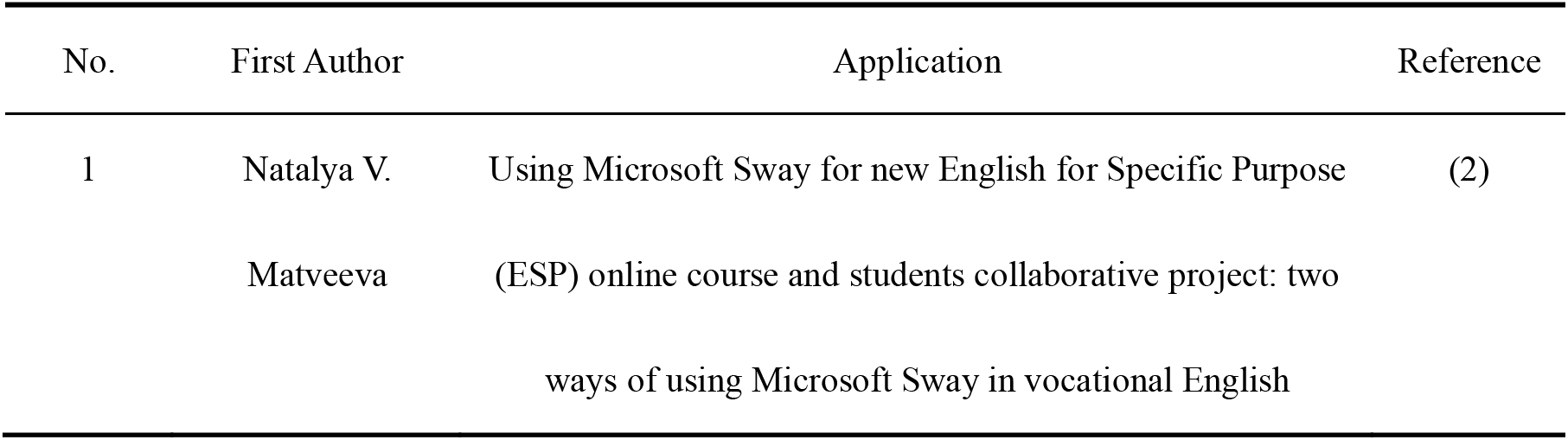
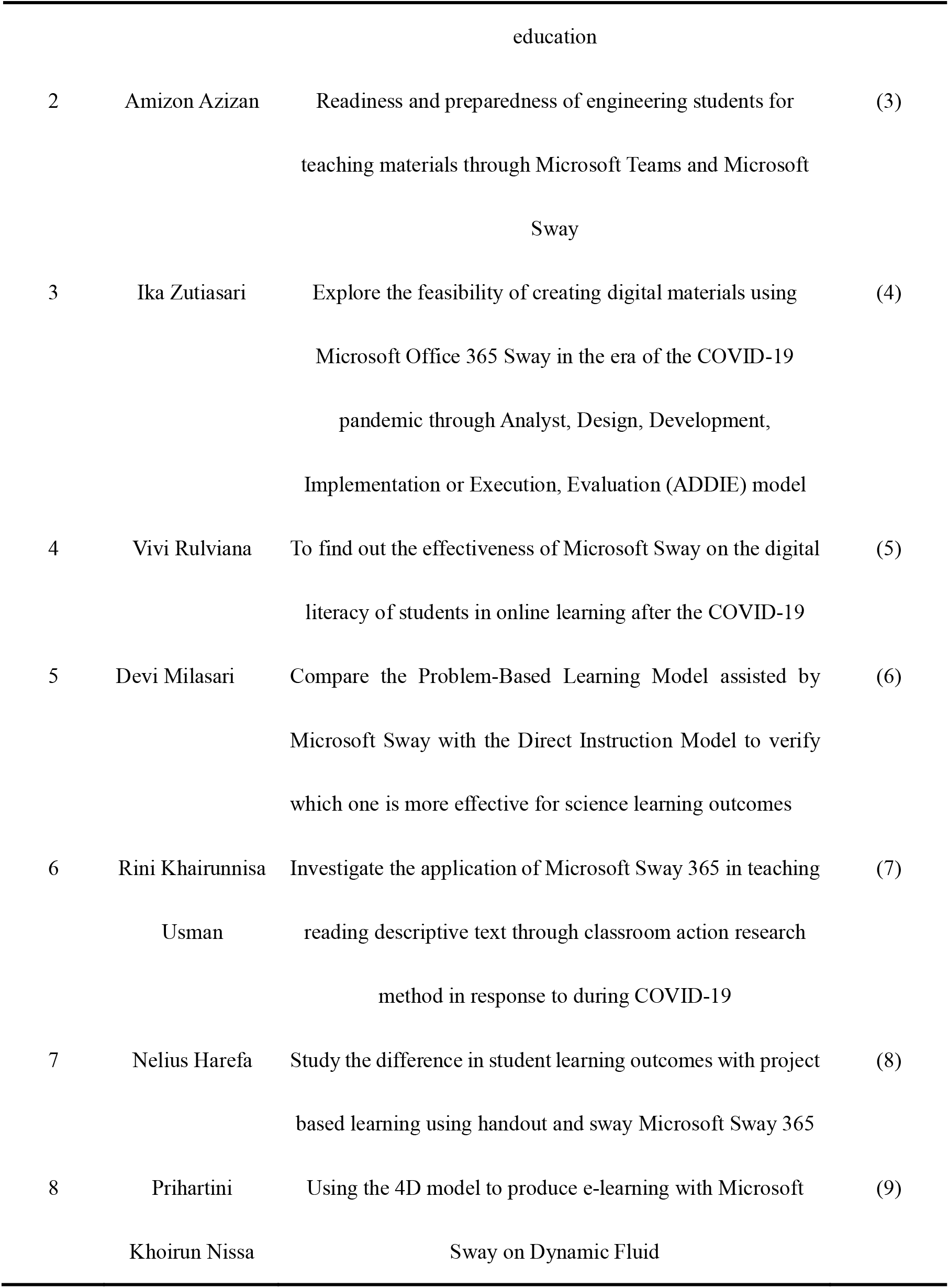
Classic studies on using Microsoft Sway in teaching research and reform.

| No. | First Author | Application | Reference |
| --- | --- | --- | --- |
| 1 | Natalya V.<br>Matveeva | Using Microsoft Sway for new English for Specific Purpose<br>(ESP) online course and students collaborative project: two<br>ways of using Microsoft Sway in vocational English education | (2) |
| 2 | Amizon Azizan | Readiness and preparedness of engineering students for teaching materials through Microsoft Teams and Microsoft Sway | (3) |
| 3 | Ika Zutiasari | Explore the feasibility of creating digital materials using Microsoft Office 365 Sway in the era of the COVID-19 pandemic through Analyst, Design, Development, Implementation or Execution, Evaluation (ADDIE) model | (4) |
| 4 | Vivi Rulviana | To find out the effectiveness of Microsoft Sway on the digital literacy of students in online learning after the COVID-19 | (5) |
| 5 | Devi Milasari | Compare the Problem-Based Learning Model assisted by Microsoft Sway with the Direct Instruction Model to verify which one is more effective for science learning outcomes | (6) |
| 6 | Rini Khairunnisa Usman | Investigate the application of Microsoft Sway 365 in teaching reading descriptive text through classroom action research method in response to during COVID-19 | (7) |
| 7 | Nelius Harefa | Study the difference in student learning outcomes with project based learning using handout and sway Microsoft Sway 365 | (8) |
| 8 | Prihartini Khoirun Nissa | Using the 4D model to produce e-learning with Microsoft Sway on Dynamic Fluid | (9) |

It should be noted that Microsoft Sway can generate a public weblink for each work generated.

Once this link is shared, the work can be viewed from any networked terminals (including mobile ones). Therefore, after completing the experimental course, students can be supported to transfer relevant original data into Microsoft PowerPoint to prepare tables and images, supplemented by text descriptions, and create links to Microsoft Sway works on this basis. Of course, students with further interest are encouraged to use professional software such as GraphPad Prism, OriginLab or R to draw images and then convert them to Microsoft Sway. After that, the monitor or class representative will summarize the Microsoft Sway links collected from each student into a single Microsoft Sway file. The summary files retain hyperlinks so they can be directed directly to individual data of students.

## Methods

Hereby, we would like to introduce a real-world case. During a pharmaceutical experiment course, salicylic acid topical ointments were prepared by students, with the indication for skin inflammation treatment.

In the experiment, three different types of ointments with different matrices were first prepared, namely (1) oil matrix ointment, (2) emulsion (O/W) matrix ointment and (3) aqueous matrix ointment. Approximately 30g of ointment were prepared totally.

1. The oil matrix ointment was prepared using the melting method. The quantity of beeswax specified in the prescription was transferred to an evaporating dish and melted in a water bath. The vegetable oil was then added slowly. Following sufficient stirring, the mixture was removed from the water bath and continued to be stirred until a viscous consistency was achieved.
2. The O/W matrix ointment was prepared via the emulsification method. Once the oil and water phases had been heated separately, the latter was added to the former, with stirring in progress, until the mixture reached the point of condensation.
3. The aqueous matrix ointment was then prepared by the grinding method, whereby methyl cellulose was placed in a mortar, along with the prescribed quantity of glycerin. As grinding occurred, the sodium benzoate aqueous solution and distilled water were slowly added, resulting in the ointment.

The combination of salicylic acid and ferric chloride (FeCl_3_) results in the reduction of iron ions in ferric chloride to ferrous iron by the hydroxyl group in salicylic acid, while the salicylic acid itself undergoes oxidation to phenol. The electrons released by the reaction will render the reaction mixture more unstable, ultimately leading to the production of a strong absorption band that exhibits a purple colour phenomenon(10, 11). Consequently, by measuring the height of the colour change interval in the agar gel containing ferric chloride over a specified period of time, the drug release rate of salicylic acid ointment can be determined.

The three salicylic acid ointments, prepared with different matrices, were placed on the agar gel containing ferric chloride, ensuring close contact and sealing with a plastic wrap. The height of the colour-developing area was measured at regular intervals to ascertain the rate of drug release.

The recorded data was imported into Origin, and a line graph was constructed of the square of the drug diffusion height versus the diffusion time to obtain the drug release curve. The line graph and table graph were exported, and the images were placed in Sway, respectively. Text descriptions were entered to export the Sway hyperlinks, and its link is https://sway.cloud.microsoft/GdI7QOVdYVrNrkpX?ref=Link.

## Results

The relevant results are presented in Figure 1 and Figure 2.

**Figure 1.**
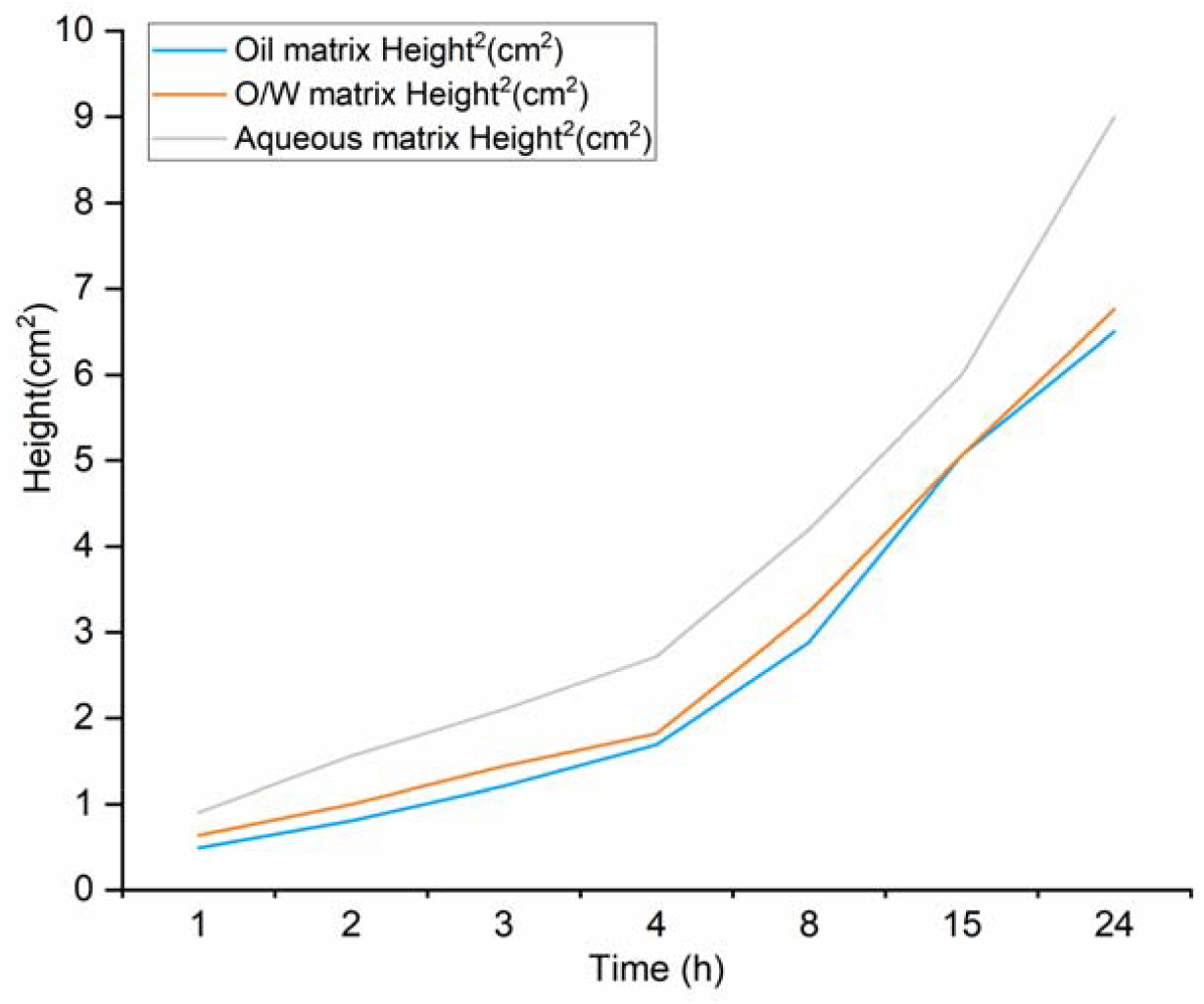
Drawing through Origin: A case on drug release from topical formulations

**Figure 2.**
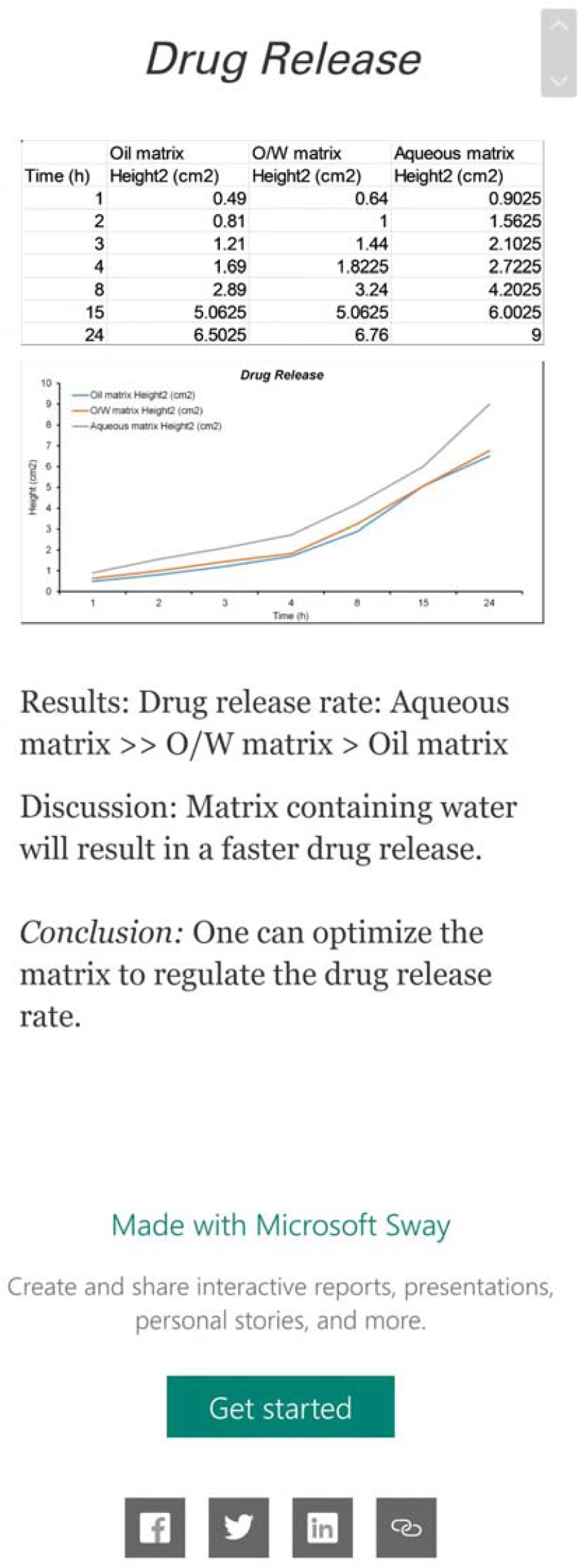
Microsoft Sway Snapshot from mobile terminal: A case on drug release from topical formulations

Experimentally, we could observe that within 24 h, the drug salicylic acid released continuously, and the release rate ranked in the sequence of aqueous matrix >> O/W matrix > oil matrix. The ramification could be explained as follows: The higher content of water in the ointment matrix, the faster drug release. It must be noted that the data should be just considered as a testing input case for subsequent Microsoft Sway processing.

A screencapture of Microsoft Sway is shown in Figure 2. Evidently, the illustration effect was acceptable. One can readily obtain and save the original data and generated charts, and interpret the discussion and conclusion. As the information could be easily shared and distributed through internet, the portability was good. This case can be used as a proof-of-concept, indicating that Microsoft Sway is capable of fulfilling the potential role of a data repository. Meanwhile, Microsoft Sway works can also be considered to incorporate into the students’ assessment process.

## Discussion

We believe that employing Microsoft Sway as a repository for data generated in pharmaceutics experimental courses has the following advantages:

- The software is simple to operate, especially without coding;
- Publicly accessible network links can be created, which can also be accessed by devices such as mobile phones and pads, and used as a portable digital dataset;
- Educators and learners are allowed to browse this repository at any time to facilitate the application prospects of mining relevant data.

In the preliminary practice of experimental pharmacy courses, we have basically confirmed the above-mentioned advantages. In addition, since Microsoft Sway actually has no specific data format requirements, it has the potential to be applied in other experimental courses, for instance, medicinal chemistry, pharmacology and pharmaceutical analysis, and can even be extended to the experimental courses in other clinical and biomedical sciences.

Of course, this strategy is still at the theoretical framework stage, and we hope to report and share it with readers as soon as possible. However, its specific outcome in improving education performance remains to be further confirmed. We will subsequently set up our own control cohort, launch prospective and retrospective studies, and conduct statistical analysis to scientifically clarify its positive outcome(12).

## Conclusion

In summary, we believe that Microsoft Sway is a great promising repository for data generated in pharmaceutics experimental courses. We have presented some relevant information from classic studies and enumerated its possible application scenarios, that is, the preparation of salicylic acid ointment and its determination of drug rate. Three salicylic acid ointments with different matrices were prepared for the experiment. The height of the colour development area was measured at regular intervals after loading them on the top of agar gel containing ferric chloride in order to obtain the drug release rate. The relevant experimental data was then imported into Origin to create a line graph. The image was subsequently imported into Microsoft Sway, accompanied by a text description. Finally, a hyperlink was generated for convenient access to the data at any time. Microsoft Sway has unique advantages such as simple operation, convenient access, and easy-to-use connection. Although it is still in the theoretical framework stage and needs to be further evaluated, its application potential in medicine-related experimental courses is beyond doubt.

